# The high infectivity of the SARS-CoV-2 Omicron variant is associated with an exclusive S477N spike receptor-binding domain mutation

**DOI:** 10.1101/2023.09.11.557161

**Authors:** Jadson C. Santos, Elvira R. Tamarozzi, Mariangela Dametto, Rodrigo Bonacin, Eduardo A. Donadi, Geraldo Aleixo Passos

## Abstract

The spike glycoprotein receptor-binding domain (RBD) of SARS-CoV-2 facilitates viral binding to the ACE2 receptor and mediates viral infectivity. The Delta and Omicron variants of concern are the most infectious strains, presenting mutated amino acid residues in their spike RBD. The Omicron variant quickly dominated the COVID-19 pandemic, indicating its greater spreadability. Omicron’s spreading might be associated with mutational substitutions at spike RBD residues. We employed *in silico* molecular dynamics (MD) simulation of the spike RBD-ACE2 interaction to compare the impact of specific mutations of the Delta and Omicron variants. The MD of the spike-ACE2 interaction showed the following: i) the amino acid profile involved in the spike-ACE2 interaction differs between Delta and Omicron; ii) the Omicron variant establishes several additional interactions, highlighting the spike RBD (S477), which is a flexible mutational residue. Since the S477N mutation is exclusive to Omicron, which may initiate binding with ACE2, the increased infectivity of Omicron might be associated not only with a mutated RBD but also with unmutated (e.g., G476 and L492) residues, initiating binding due to the influence of the N477 mutation. Compared to previous variants, Omicron’s N477 residue represents a novelty within the spike-ACE2 interaction dynamics interface.

## Introduction

Enveloped RNA viruses, such as SARS-CoV-2, exhibit higher mutation rates than DNA viruses (1). A typical SARS-CoV-2 genome accumulates only two single-nucleotide mutations per month, i.e., less than other RNA viruses, such as HIV-1 or influenza (2). The fast spread of SARS-CoV-2 in humans is driving viral molecular evolution mainly among unvaccinated persons (3-6) and in patients with immunocompromising conditions (7-9).

Although most mutations found in SARS-CoV-2 are neutral, some may affect viral replication or infectivity (1,10-13. Since the earliest viral isolates up to the last SARS-CoV-2 structure (14), several nucleotide changes in the viral genome have been reported. As of the conclusion of this manuscript in late August 2023, one of the most recent Omicron variants of concern (VOCs) reported (HCoV-19/Singapore/R13MR39/2023, clade 21 L), identified in Asia, has accumulated nucleotide changes, reversions to root, gaps, and amino acid changes in the Spike (S1) mRNA/protein (nextstrain.org/ncov/gisaid/global/6 m). Most of the variability found in SARS-CoV-2 isolates is observed at the spike receptor-binding domain (RBD), a portion of the glycoprotein that mediates viral attachment to the angiotensin-converting enzyme 2 (ACE2) receptor on the surface of human cells (15-19).

As previously reported (20), emergence of the SARS-CoV-2 alpha (B.1.1.7) variant as early as December 2020 (21, 22) represents an example, among several others, of rapid viral molecular evolution (23). The Alpha (B.1.1.7) phenotype has also attracted attention due to its high transmissibility among humans (21, 22), accumulating 17 mutations in its genome including 8 in the gene encoding the spike protein, which is present on the virus surface (22). In particular, the N501Y mutation in the spike gene covers one of the six key contact amino acid residues within the spike RBD of Alpha (B.1.1.7) (22). Considering that the N501 mutation increases binding affinity to human ACE2 (9), mapping of the amino acid residues that make contact with human ACE2 may reveal desirable targets for developing vaccines against or for antibody-based therapies for COVID-19 (24, 25).

At present, the Wuhan original lineage and five VOCs are present in the WHO SARS-CoV-2 variant dataset, i.e., Alpha (B.1.1.7), Beta (B.1.351), Gamma (P.1), Delta (B.1.617.2), and Omicron (B.1.1.529), the last of which was identified in November 2021 (26) in South Africa and recently with its new sub-lineage EG.5 emerged in early 2023 (https://www.gisaid.org/). Genome sequencing of the Omicron variant has shown 35 nonsynonymous mutations in spike’s amino acid residues, 15 of which are in the receptor-binding domain (RBD) and 10 in the receptor-binding motif (RBM); these mutations may explain its high rate of spread (27). Therefore, the impact of SARS-CoV-2 mutation on spike RBD affinity for the ACE2 receptor is crucial for understanding the molecular basis of the infectivity/transmissibility of SARS-CoV-2 VOCs.

Considering that the unmutated RBD and the N501Y mutation exhibit distinct interactions with ACE2, we previously hypothesized that N501Y increases the strength of the interaction between the spike RBD and the ACE2 receptor. Indeed, we demonstrated the plausibility of this idea through *in silico* mutagenesis and spike interaction analyses (20). As new infective variants of epidemiological importance have been described, such as the Delta and Omicron variants, in this study, we sought to determine whether the variability of the amino acid residues found in these two SARS-CoV-2 VOCs interferes with the interaction at the spike-ACE2 interface and whether the interaction stability between the spike-ACE2 interfaces differs between these VOCs. We observed that specific amino acid residues between the two spike proteins are associated with the infectivity/transmissibility of these VOCs, particularly the new Omicron variant S477N mutation, which may be associated with its increased infectivity.

## Results

The free energy of the RBD-ACE2 complexes of the Delta (B.1.617.2) and Omicron (B.1.1.529) variants showed similar values (−4.2186325e+06 and -4.0165590e+06 kJ/mol, respectively), demonstrating that, at least in quantitative terms, the complexes behave similarly.

### Root mean square deviation (RMSD) molecular dynamics

We observed no significant difference in the stability of the interaction for the SARS-CoV-2 spike receptor-binding domain (RBD) with the ACE2 receptor. However, at the final dynamics time, the Delta variant was slightly less stable than the Omicron variant (Figure 1).

**Figure 1.**
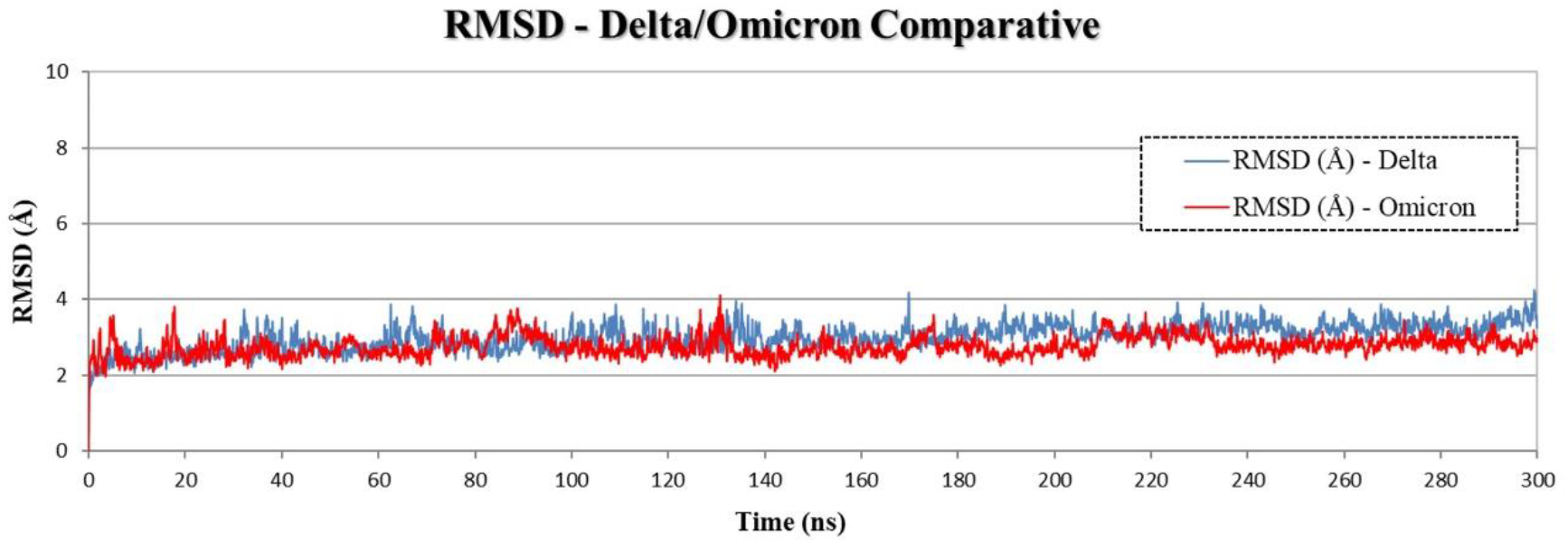
Molecular dynamics simulation between SARS-CoV-2 Spike receptor binding domain and ACE2 receptor through RMSD comparing Delta (blue) and Omicron (red) variants of concern. Delta variant exhibited lesser stability of spike RBD-ACE2 interaction compared to Omicron variant at 240-300 ns dynamics time.

### B-factor calculation

Our b-factor analysis for the S477N mutation region (Figure 2) showed more significant movement in the region of amino acids 477 to 484 of the Spike RBS of the Delta variant than that of Omicron. The loop formed by these amino acids is essential in interacting with the ACE2 receptor. This difference was induced by the S477N mutation, promoting less flexibility and greater structural thermostability to the loop in the Omicron variant. Thus, increased capacity and effectiveness in the RBD-ACE2 interaction might result in Omicron’s higher infectivity.

**Figure 2.**
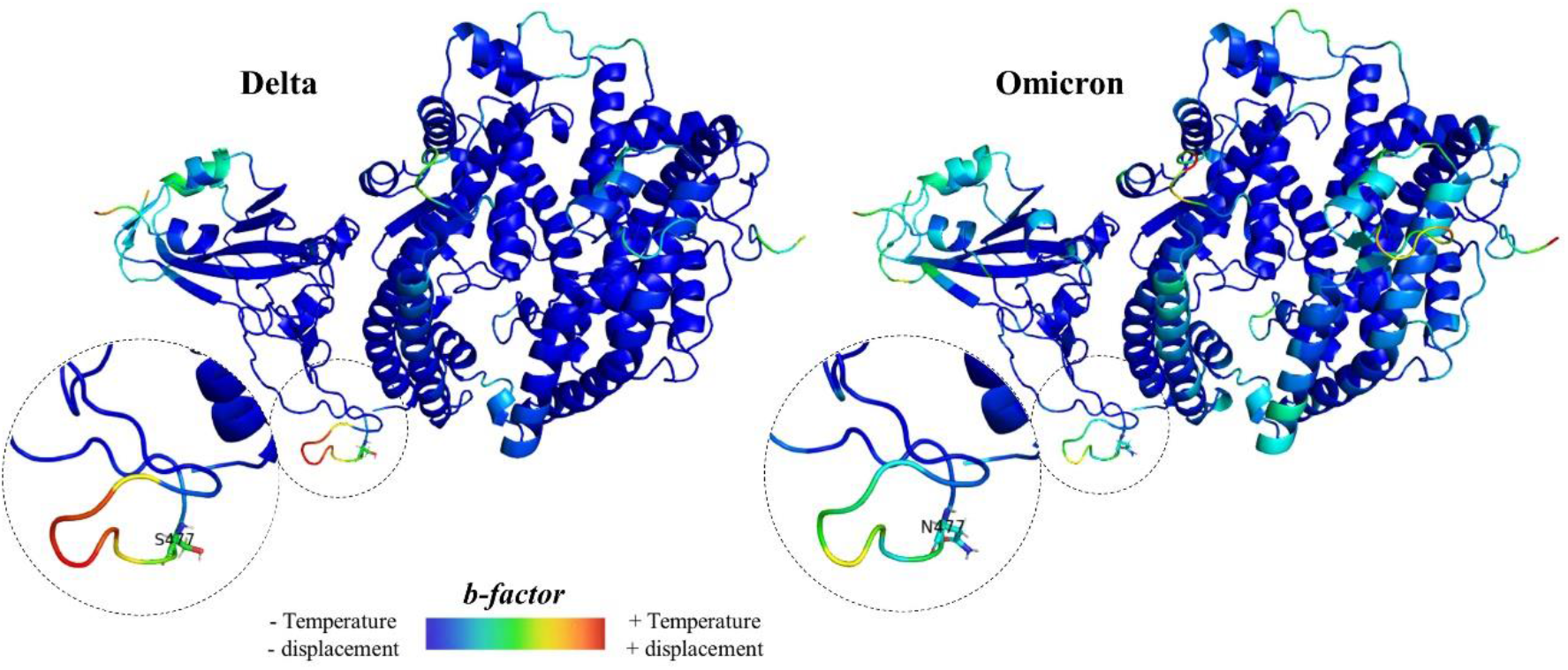
The B-factor analysis of the Omicron’s S477N mutation region. This analysis showed more significant movement in the Delta to Omicron variant’s amino acid segment 477 to 484 Spike RBS. The noted variance was caused by the S477N mutation, which conferred the Omicron variant with reduced flexibility and enhanced thermal stability in its loop structure.

### Root mean square fluctuation (RMSF) molecular dynamics

We also explored the dynamics of residues in terms of root mean square fluctuation (RMSF) analysis, which can reveal the areas of the RBD-ACE2 complex, which is more mobile. The RMSF results showed similar fluctuations, with different magnitudes for the Delta and Omicron systems, i.e., compared to Omicron variant, the Delta variant displays a region with higher mobility from residues 349 to 529 (Figure 3).

**Figure 3.**
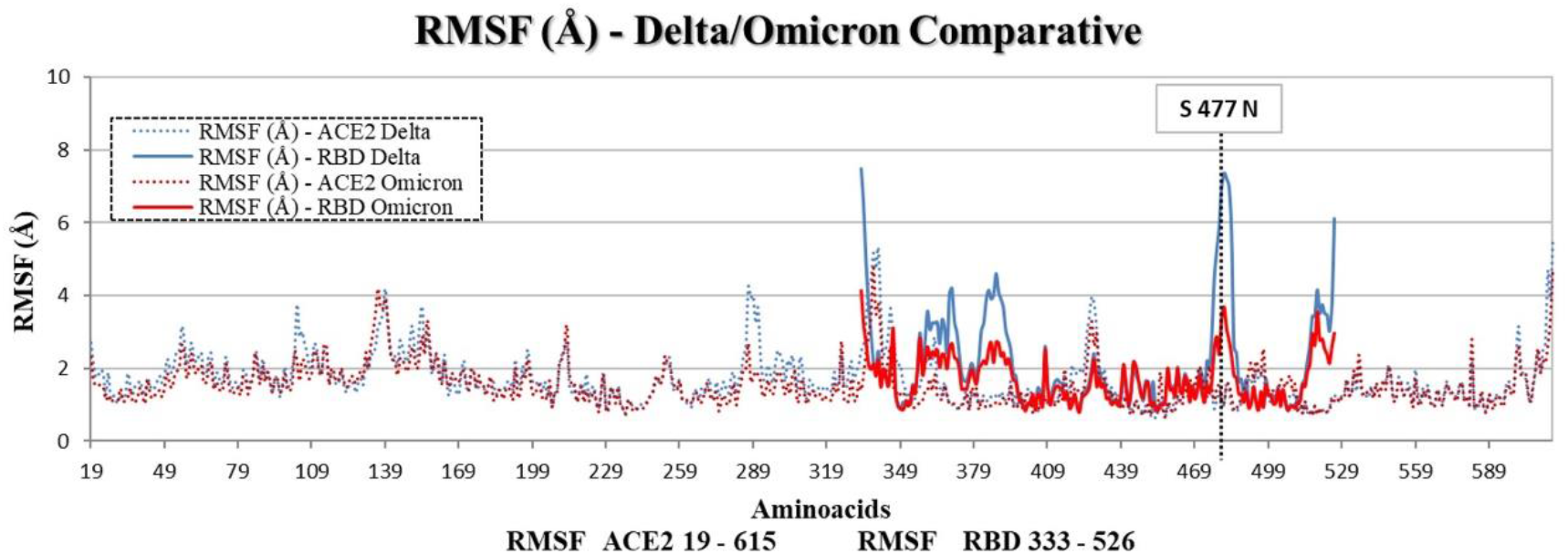
Molecular dynamics simulation between SARS-CoV-2 Spike receptor binding domain and ACE2 receptor through RMSF comparing Delta variant (blue) and Omicron variant (red) variants of concern. The dashed vertical line indicates the position of the S477N mutation. The RMSF findings demonstrated analogous fluctuations, albeit with distinct magnitudes for Delta and Omicron systems. A region exhibiting higher mobility from residues 349 to 529, was more evident in the Delta variant.

### Interaction analysis

After the simulation time of 300 ns, comparisons of the spike RBD-ACE2 interaction interface of the two VOCs showed quantitative similarity regarding the total number of contacts.

Regarding unique contacts, the Delta and Omicron variant complexes showed a profile in which the latter expands the interface laterally (residues L492, K458, G476, and N477) (Figure 4). In contrast, the Delta variant maintains most of the unique interface residues at other interaction regions.

**Figure 4.**
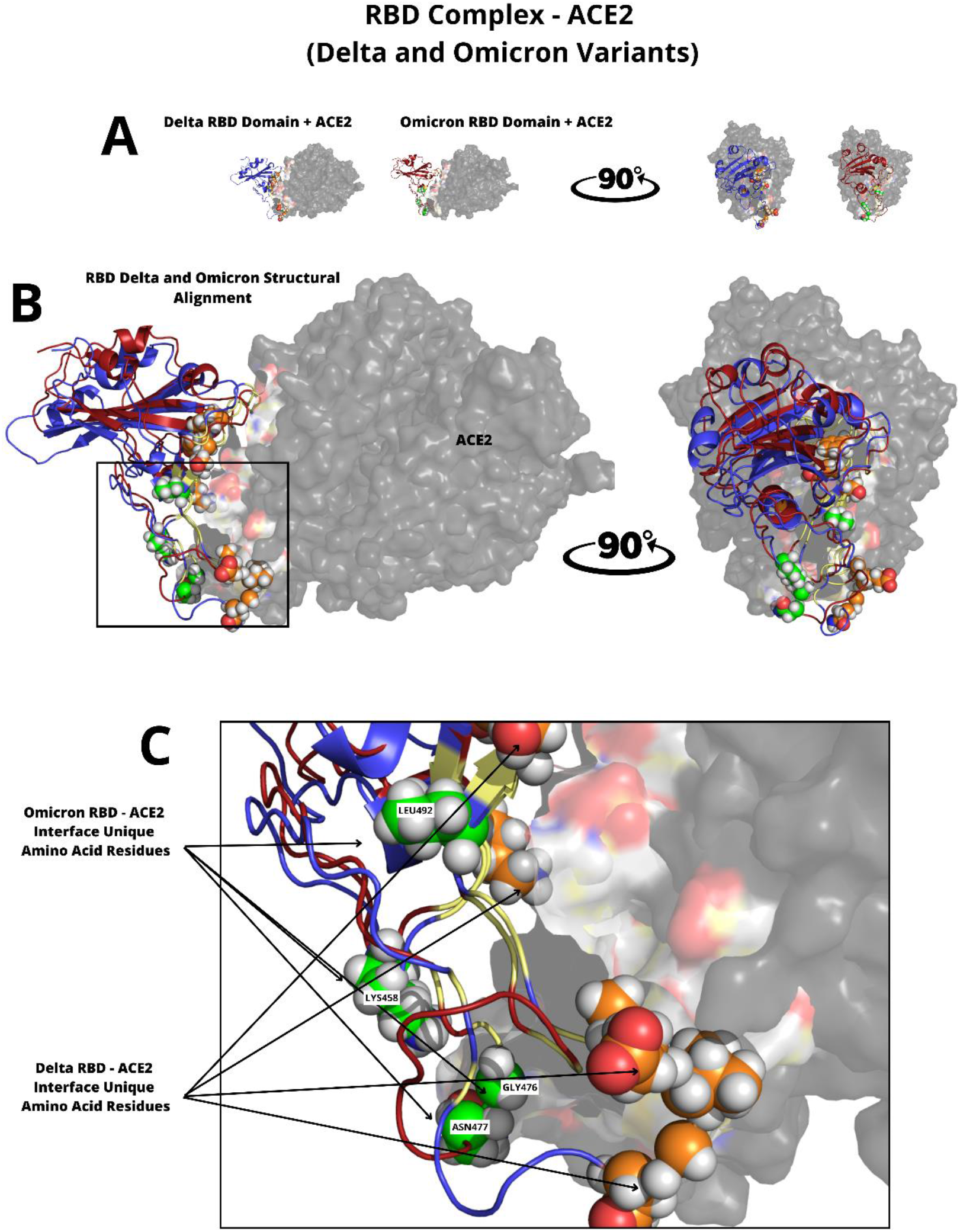
Identifying specific amino acid residues present in the lateral expansion of the Omicron spike RBD-ACE2 interface. The unique RBD amino acid residues in the RBD-ACE2 interface are represented as spheres. A) Delta (blue) or Omicron (red) variant RBD-ACE2 complexes are shown with different rotation angles. B) The Delta or Omicron variant complete RBD-ACE2 complexes are shown in structural alignment and rotated by 90°. C) The particular Omicron RBD-ACE2 interaction resulting from the S477N mutation (Omicron in green and Delta in orange).

After identifying this lateral expansion of the interaction interface between the RBD domain of the Omicron variant and human ACE2, we sought to determine which specific residues are present in this region. We identified that a single mutation at residue S477N of the RBD domain of Omicron variant creates a new contact point that involves not only the mutant residue but also two other nonmutated residues, G476 and L492 (Figure 4).

## Discussion

Considering that the two most recent SARS-CoV-2 variants, Delta and Omicron, do present amino acid substitutions at the spike glycoprotein receptor binding domain (RBD) that may increase the interaction of the virus with the cellular receptor ACE2 (Spike-ACE2 interaction) (15-19), we studied the molecular mechanisms of these interactions that may be advantageous for the virus, increasing its infectivity.

The SARS-CoV-2 Omicron B.1.1.529 variant was identified in late November 2021, as noted by the WHO (28). It has since dominated the COVID-19 pandemic as a highly spreading/infective strain, despite being considered less pathogenic than the Delta variant (29-31). In a previous study (20), we hypothesized that the rapid spread of SARS-CoV-2 variants might be associated with mutational substitutions in spike RBD residues. Here, we use *in silico* molecular dynamics simulations of the spike RBD-ACE2 interaction complex to compare the Delta and Omicron variants, evaluating amino acid substitutions that might be associated with the rapid spread of Omicron variant.

By comparing the spike RBD sequence between the two variants, the following substitution mutations have been described: i) Delta variant (L452R and T478K) and ii) Omicron variant (G339D, S371L, S373P, S375F, K417N, N440K, G446S, S477N, T478K, E484A, Q493K, G496S, Q498R, N501Y, and Y505H) (32). These mutations might influence viral neutralization by antibodies and viral escape (33-35) or may benefit the virus by increasing its infectivity compared to the original Wuhan strain (32).

Investigation of the spike RBD-ACE2 binding affinity is crucial to better understand viral infectivity. However, this aspect is still unresolved and somewhat controversial in the literature, as illustrated by several lines of evidence, including from *in vitro* and *in silico* studies. An increase in the strength of the interaction between spike and ACE2 has been reported as a result of spike mutations in combination as an escape mechanism of the humoral immune system response (36, 37). Compared to the interaction force of the Wuhan original strain, i) the Alpha variant presents an approximately 10-fold increase due to acquisition of the RBD N501Y mutation (34), ii) the Beta variant exhibits a 5-fold increase in the interaction strength of the RBD-ACE2 complex (36), and iii) the Delta variant shows a 2-fold increase (38). Molecular dynamics simulations indicate that the Omicron variant presents binding affinity to ACE2 that is similar to the Wuhan original strain; however, it exhibits weaker binding affinity compared to the Delta variant (39). Using ELISA, these authors found that ACE2 binding affinities with Omicron variant and the original Wuhan VOCs are similar.

Here, we report that Delta and Omicron variants have similar molecular dynamics simulations. Therefore, total free energy determinations and stability of the spike RBD-ACE2 complexes may not explain the observed differences in affinity between these proteins or increased viral infectivity when comparing these variants. We and others (34) have found that the Omicron variant harbor the spike RDB N501Y mutation (that was observed in the Alpha and Delta VOCs but not in the Wuhan strain), which may contribute to increased affinity between RBD-ACE2 compared to the original Wuhan strain. Therefore, the N501Y mutation, which is in direct contact with ACE2, would not explain Omicron’s exclusive characteristic of high infectivity.

Mutations in residues that are not in direct contact with ACE2 may impact the conformational dynamics of the RBD domain to promote more efficient binding of Delta and Omicron to ACE2 (40). Among the mutations present in the Omicron variant, the functional impacts of some have been reported, such as K417N, S447N, E484A, and Q493R, which may contribute to immune escape responses (41), and N501Y, which may contribute to virus infectivity (20).

Our analysis of the spike S1 portion between residues 475 to 485, which is highly flexible, showed that position S477 is the most flexible and the most exchangeable residue among SARS-CoV-2 RBD mutants. The prediction that the S477G and S477N mutations increase spike-ACE2 binding was published months before the emergence of the Omicron variant in late 2021 (42). In recent work (43, 44), the authors use computational analysis, evaluated position 477 of the Spike protein, and concluded that the S477N mutation found in the Omicron variant increases the interaction with the ACE2 receptor.

In this study, we observed that the S477N mutation could turn nonmutated residues (G476 and L492) into part of the RBD-ACE2 interface. These contacts, in combination with the S477N mutation, may strengthen the RBD-ACE2 interaction. The S477N mutation is present in the Omicron variant but not in any other previous VOC, playing a central role in interacting with ACE2 and further promoting expansion of the RBD-ACE2 interface. In 2020, before the Delta and Omicron variants appeared, researchers used a quantitative deep mutational scanning approach to experimentally measure how all possible RBD amino acid mutations might affect RBD-ACE2 binding affinity (9). Among the analyzed mutations, they found that the S477N mutation positively affect expression of a folded RBD protein and its affinity for ACE2, two critical factors for viral fitness (9). Our molecular dynamics studies showed that the S447N mutation is essential for interacting with human ACE2, recruiting other residues at the interface. In this context, our data validate the prediction (9) and expand the importance of this mutation to the RBD-ACE2 interface. The importance of the mutated residue (N477) in the Omicron variant was further reported (43), whose authors demonstrated that the residue begins to interact due to hydrogen bonding interactions. In our study, we show how nonmutated residues (G476 and L492) close to the S477N mutation begin to interact under the influence of this mutation. Others (42) also share this view, who concluded their computational research by stating that although some favorable hotspot residues for ACE2 binding are disrupted in the Omicron RBD, new binding interactions are promoted in this variant.

Using a combinatorial approach, researchers (37) identified that some combinations of mutations have a potential role in increasing affinity of the spike protein toward ACE2. Among these combinations, S477N/E484K presents resistance against neutralizing antibodies in addition to increased affinity for ACE2. As reported in our study, position 484 is present in the central region of the RBD-ACE2 interface in both Delta and Omicron. This region is the target of most neutralizing antibodies (41); however, position 477 is located in the interface’s lateral region. As we have shown, as the S477N mutation is exclusive to Omicron, this residue is not involved at the interface in the Delta variant. Therefore, it is possible that the lateral region of the interface, in which the S477N mutation is located, plays a central role in affinity by binding this residue and sequestering residues G476 and L492 to the interface. In contrast, central region mutations are more critical for viral escape.

## Methods

### Spike RBD-ACE2 interface analysis and in silico *mutagenesis*

#### Molecular dynamics simulation

We performed a 300 ns molecular dynamics simulation of the three-dimensional structure of the spike RBD domain from SARS-CoV-2 Delta (B.1.617.2) and Omicron (B.1.1.529) in complex with the ACE2 extracellular domain. To simulate the mutations present in the spike RBD domain of the variants, we used an RBD-ACE2 complex structure available in Protein Data Bank (45) (code 6m0j) using the Wuhan spike RBD domain.

The PyMOL software (46, 47) mutagenesis tool was used to perform amino acid substitutions (mutations). GROMACS software (38, 48) was used for molecular dynamics simulation, implementing the CHARMM36m force field (50). A previous step of energy minimization was applied to the protein/solvent system submitted to molecular dynamics simulation, and afterward, we balanced the system using GROMACS. Next, we calculated the atomic behavior and temporal evolution of the atoms that comprise the system (trajectory). At the end of the molecular dynamics simulation time, files were generated through GROMACS. Graphs were generated using Excel to visualize the trajectory from the molecular dynamics simulation.

The three-dimensional structures obtained along the molecular dynamics trajectory were visualized using PyMOL. Root mean square deviation (RMSD) and root mean square fluctuation (RMSF) values were estimated using the α carbons (Cα) of the initial structure as a reference.

The RMSD calculates the mean square deviation between all atoms of a three-dimensional structure relative to a reference structure (51); it also calculates the mean square deviation of the fluctuation of the positions of each atom in the structure relative to a reference structure (52) over the given simulation time. The RMSF values calculated from molecular dynamics simulations measure the mobility of each amino acid around its median position in the structure and allow assessment of protein flexibility (54). The b-factor (known as the temperature factor) measures temperature-dependent atomic displacement and vibration, representing a quantitative indication of the flexibility of a protein structure (39).

#### B-factor calculations

For each molecular dynamics simulation, the b-factor was calculated using the initial structure of the total protein as a reference. The calculations were performed using GROMACS software (48, 49). The b-factor, known as the temperature factor, is a temperature-dependent measure of atomic displacement and vibration. It presents a quantitative indication of the flexibility and thermostability of a protein structure (53, 54).

## Author’s contributions

JCS: Performed definitive molecular analysis, constructed the figures, interpreted and discussed the results and wrote the manuscript; ERT: Performed definitive molecular analysis, constructed the figures, interpreted and discussed the results; MD: Performed molecular analysis at the beginning of the project to evaluate its viability; RB: Performed molecular analysis at the beginning of the project to evaluate its viability; EAD: Interpreted and discussed the definitive results and wrote the manuscript; GAP: Conceived the study, evaluated the project’s viability, raised hypothesis, interpreted and discussed the results and wrote the manuscript.

## Data availability

The datasets used and/or analyzed during the current study available from the corresponding author on reasonable request. The viruses analyzed in this work were registered with the Brazilian Genetic Resources Council, CisGen Register No. A917837.

